# Evidence that genetic drift not adaptation drives *fast-Z* and *large-Z* effects in *Ficedula* flycatchers

**DOI:** 10.1101/2023.02.08.527632

**Authors:** Madeline A. Chase, Maurine Vilcot, Carina F. Mugal

## Abstract

The sex chromosomes have been hypothesized to play a key role in driving adaptation and speciation across many taxa. The reason for this is thought to be the hemizygosity of the heteromorphic part of sex chromosomes in the heterogametic sex, which exposes recessive mutations to natural and sexual selection. The exposure of recessive beneficial mutations increases their rate of fixation on the sex chromosomes, which results in a faster rate of evolution. In addition, genetic incompatibilities between sex-linked loci are exposed faster in the genomic background of hybrids of divergent species, which makes sex chromosomes contribute disproportionately to reproductive isolation. However, in birds, which show a Z/W sex determination system, the disproportionate role of the Z-chromosome in adaptation and reproductive isolation is still debated. Instead, genetic drift has been proposed as the main driver of the so-called *fast-Z* and *large-Z* effects in birds. Here, we address this question in *Ficedula* flycatchers based on population resequencing data of six flycatcher species. Our results provide evidence for both the *fast-Z* and *large-Z* effects in *Ficedula* flycatchers and that these two phenomena are driven by genetic drift rather than positive selection. Genomic scans of selective sweeps and fixed differences in fact suggest a reduced action of positive selection on the Z-chromosome. We propose that the observed reduction in the efficacy of purifying selection on the Z-chromosome helps to establish genetic incompatibilities between Z-linked and autosomal loci, which could result in pronounced selective sweep signatures for compensatory mutations on the autosomes.

## Introduction

The number and genomic distribution of genetic loci that contribute to adaptation and reproductive isolation is a central question in speciation research. Speciation genomic studies across a wide range of taxa have revealed a heterogenous differentiation landscape along the genome, where the heterogeneity is frequently attributed to divergent selection and so-called barrier loci that build up resistance to gene flow earlier than the genomic background (Feder et al., 2012; Nosil et al., 2009; Via & West, 2008). In addition, the variability in recombination rate and thus in the intensity of linked selection are known to contribute to the heterogeneity in differentiation (Nachman & Payseur, 2012; Ravinet et al., 2017; Wolf & Ellegren, 2017). Yet, since sex chromosomes expose incompatible genetic loci in hybrids of the heterogametic sex faster, sex chromosomes are assumed to make a disproportionally large contribution to hybrid dysfunction and reproductive isolation, commonly referred to as the ‘*large-X* (or *large-Z*) effect’ (Coyne, 1984; Presgraves, 2008; R. Storchová et al., 2010). Furthermore, sex-linked genetic loci are exposed to selection more efficiently in the heterogametic sex, which increases the efficacy of selection on sex chromosomes (Avery, 1984). Provided that beneficial mutations are on average recessive, this may lead to faster evolution of the X (or Z) chromosome, commonly referred to as the ‘*fast-X* (or *fast-Z*) effect’ (Charlesworth et al., 1987). In line with these hypotheses, genomic studies have revealed elevated differentiation levels on sex chromosomes compared to autosomes (Presgraves, 2018). However, elevated differentiation on sex chromosomes does not necessarily reflect a disproportionate contribution of sex chromosomes to adaptation and reproductive isolation (Coyne, 2018; Presgraves, 2018). The reasons for elevated differentiation on sex chromosomes can in fact be manifold.

The sex chromosomes spend unequal times in males and females, and are therefore differently affected by sex-specific selection mechanisms (Charlesworth et al., 1987; Rice, 1984) and demography (Pool & Nielsen, 2007). Moreover, sex chromosomes and autosomes have different effective population sizes (*N*_e_) (Vicoso & Charlesworth, 2009), mutation rate (Ellegren, 2007; Hedrick, 2007; Kirkpatrick & Hall, 2004), and recombination rate (Hedrick, 2007), which can all contribute to differences in the genomic differentiation landscape among sex chromosomes and autosomes. Since birds have a female heterogametic sex determination system (ZW females and ZZ males), we will in the following discuss these differences from the angle of a ZW sex determination system. Similar arguments apply to an XY sex determination system but with switching the sexes (Irwin, 2018).

Given an equal proportion of reproducing females and males in the population, the Z-chromosome versus autosome (Z:A) ratio of *N*_e_ is 3/4 (Vicoso & Charlesworth, 2009), but can range between 9/16 to 9/8 if differences in reproductive variance between sexes are present. These differences in *N*_e_ among the Z-chromosome and autosomes will naturally impact the selection-drift balance and for commonly observed Z:A ratios < 1 lead to a lower efficacy of selection on the Z-chromosome. Indeed, ample evidence suggests that the *fast-Z* effect in birds is more likely a result of less efficient purging of deleterious mutations due to a higher impact of genetic drift rather than a result of more efficient positive selection in the heterogametic females (Hayes et al., 2020; Mank et al., 2010; Wang et al., 2014). Besides affecting the efficacy of selection, a lower *N*_e_ for the Z-chromosome also increases the speed of lineage sorting (Presgraves, 2018; Wolf & Ellegren, 2017), which results in faster genomic differentiation (or higher *F*_ST_) on the Z-chromosome than the autosomes. Genomic signatures of a *large-Z* effect that purely rely on elevated genomic differentiation therefore do not necessarily need to invoke hybrid incompatibilities. Moreover, mutation rate is generally found to be male-biased in birds (Axelsson et al., 2004; Wang et al., 2014), which increases genomic differentiation and could be another confounding factor of signatures of *fast-Z* or *large-Z* effects. Finally, since the heteromorphic parts of sex chromosomes only recombine in the homogametic sex, the Z-chromosome is observed to show lower rates of recombination compared to autosomes across birds (Wang et al., 2014). On the one hand, a lower recombination rate can result in a reduced efficacy of selection, and thus further affect the selection-drift balance on sex chromosomes. In addition, genome-wide signatures of introgression have been shown to positively correlate with recombination rate, with lower levels of introgression coinciding with low recombination (Martin et al., 2019; Schumer et al., 2018). Consequently, this mechanism could mimic a *large-Z* effect if not properly accounted for. In light of all these possible confounding factors, the investigation of the role of sex chromosomes in adaptation and reproductive isolation therefore requires a thorough evaluation.

Here, we address this question in *Ficedula* flycatchers, which are an important avian speciation model (Qvarnström et al., 2010; G.-P. Sætre & Sæther, 2010). In particular, the naturally hybridizing collared flycatcher (*Ficedula albicollis*) and pied flycatcher (*F. hypoleuca*) have been intensively studied in the context of speciation, and the Z-chromosome has been proposed a hotspot for reproductive isolation and adaptive speciation (Borge et al., 2005; Sæther et al., 2007; Sætre et al., 2003). However, earlier studies on the role of the Z-chromosome in the speciation process of collared and pied flycatchers have primarily been based on a few markers. More recent genome-wide approaches, on the other hand, are each limited to one type of genomic signature and do not address the role of the Z-chromosome in speciation (Nadachowska-Brzyska et al., 2019; Nater et al., 2015). To provide a comprehensive evaluation of the *fast-Z* and *large-Z* effects in *Ficedula* flycatchers, we therefore take advantage of a rich amount of genomic resources comprising of population resequencing data of six species: three populations of collared flycatcher, three populations of pied flycatcher, one population of Atlas flycatcher (*F. speculigera*), one population of red-breasted flycatcher (*F. parva*), one population of taiga flycatcher (*F. albicilla*), and one individual of snowy-browed flycatcher (*F. hyperythra*). With this dataset, we assess evidence for the *fast-Z* and *large-Z* effects through genomic signatures of direct selection, linked selection and gene flow in light of possible confounding factors.

## Methods

### Variant calling and filtering

We compiled a data set of single nucleotide variant (SNV) calls for 187 samples of four *Ficedula* flycatcher species, i.e. collared flycatcher, pied flycatcher, red-breasted flycatcher and taiga flycatcher, and one outgroup species, snowy-browed flycatcher, for the autosomes and the Z chromosome. Here, SNV calls represent single nucleotide polymorphisms (SNPs) within species and single nucleotide differences among species. Reads for all species were mapped to the collared flycatcher reference genome FicAlb1.5 (Kawakami et al., 2014). Variant calls for scaffolds belonging to the autosomes were retrieved from (Chase & Mugal, 2022). Variant calling for Z-chromosome scaffolds was performed within the present study, where we followed previously described methods (Chase & Mugal, 2022), with additional filtration steps applicable to the Z-chromosome. Briefly, genotyping was performed using GATK v.4.1 Haplotype Caller for all individuals separately, followed by joint genotyping with GenotypeGVCFs. Genotyping was performed using the flag *--all-sites* to genotype both polymorphic and monomorphic positions. For the Z-chromosome scaffolds we applied a filter of GQ >30 for male samples and GQ >15 for female samples, since females have only one copy of the Z-chromosome in birds. Additionally, we removed any sites with heterozygous genotypes in females after applying genotype filters. Finally, after performing all filters, we removed sites with more than 10% missing data in any of the four species. Our final dataset included 51,424,863 SNVs within a set of 566,724,393 callable sites in total on the autosomes from (Chase & Mugal, 2022), combined with 2,662,105 SNVs within a set of 30,171,840 callable sites in total on the Z-chromosome.

### Estimates of the Z/A ratio of effective population size

We computed four estimates of the ratio of effective population size (*N*_e_) on the Z-chromosome compared to the autosomes. First, we calculated the ratio of nucleotide diversity π_Z_/π_A_ across the entire assembled autosomes and the Z-chromosome. We calculated π for the autosomes and the Z-chromosome following the equation:

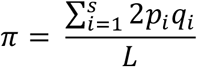

where *p*_i_ and *q*_i_ represent the allele frequencies at site *i* in *s* variable sites. We then obtained the per site measure of π by dividing by the total number of callable sites, *L*, from the all-sites vcf. We estimated π using either all SNPs for both the autosomes and the Z-chromosome, or only GC-conservative SNPs (Strong-to-Strong: S-to-S and Weak-to-Weak: W-to-W) in order to account for GC-biased gene conversion (gBGC) (Bolívar et al., 2018). To obtain confidence intervals, we randomly resampled variable sites with replacement, for 1000 bootstrap replicates. Second, we calculated the ratio of nucleotide diversity π_Z_/π_A_ after masking sites potentially affected by linked selection. For this purpose, we masked sites in the reference genome overlapping with both exons (Ensembl version 104; Uebbing et al., 2016) and conserved noncoding elements with a minimum size of 100bp (CNEs; Craig et al., 2018), and an additional 1kb flanking region on both sides of the exons and CNEs.

Additionally, we computed historical variation in *N*_e_ separately for the Z-chromosome and the autosomes using the Pairwise Sequentially Markovian Coalescent (PSMC) model implemented in the PSMC software (Li & Durbin, 2011). We performed PSMC estimation for one individual of each of the four species (Table S1), which were chosen to minimize differences in mean coverage between sequences as this can bias estimates of *N*_e_ (Nadachowska-Brzyska et al., 2016). Sites with a read depth below 10 were excluded, and we masked sites with more than twice the average read depth across the genome. Blocks of 100bp containing more than 20% missing data were excluded.

To run PSMC, we followed (Nadachowska-Brzyska et al., 2016), setting the parameters to -p “4+30*2+4+6+10”, -t5 and -r1. We performed 100 bootstrap replicates by splitting chromosome sequences into segments with ‘splitfa’ and randomly sampling segments with replacement. *N*_e_ estimates were re-scaled with the ‘psmc.results’ function from (Liu & Hansen, 2017) customized by Leroy et al. (2021), using a generation time of 2 years and a mutation rate of 4.6∙10^−9^ per site per generation (Smeds et al., 2016). We then estimated the ratio of the harmonic mean *N*_e_. For each species, we took the average *N*_e_ estimates from PSMC in 1000-year discrete time steps from the most recent time up until 1mya, separately for the Z-chromosome and the autosomes. Finally, we estimated 95% confidence intervals based on the harmonic mean *N*_e_ for each of the 100 bootstraps.

To compute estimates of the ratio of *N*_e_ on the Z-chromosome versus the autosomes either from the ratio of diversity or PSMC estimates, we followed (Irwin, 2018) and corrected the Z-chromosome values for male-biased mutation rate by dividing by 1.1.

### Ancestral sequence reconstruction

We polarized SNPs in the four ingroup *Ficedula* flycatcher species using snowy-browed flycatcher as an outgroup, which were then used for selective sweep detection (see below). For this purpose, we combined collared and pied flycatcher samples to form a second outgroup to polarize SNVs for red-breasted and taiga flycatchers, and vice versa. An allele was identified as ancestral when any two of the three groups (snowy-browed flycatcher, collared and pied flycatcher, or red-breasted and taiga flycatcher) were fixed for the same allele. Using this approach, we were able to polarize 49,121,805 SNVs on the autosomes and 2,561,370 SNVs on the Z-chromosome.

With the polarized sites we then reconstructed the ancestral sequence from the collared flycatcher reference genome (version FicAlb1.5). We first masked sites that were not genotyped based on the allsites VCF. Then, we masked variable sites that were unable to be polarized, as their ancestral state is equivocal. Finally, we replaced the collared reference allele with the ancestral allele.

### Estimates of selection in protein-coding regions

We estimated selection in protein-coding sequences on the autosomes and the Z-chromosome based on the ratio of non-synonymous over synonymous nucleotide diversity (*d*_N_/*d*_S_), the ratio of non-synonymous over synonymous nucleotide divergence (*d*_N_/*d*_S_), and the adaptive rate of evolution (ω_a_). We estimated *d*_N_/*d*_S_ for all four species by identifying 0-fold and 4-fold degenerate sites, from coding sequences for the ancestral genome reconstruction described above. We then subset the polymorphic sites to only GC-conservative polymorphisms (Strong-to-Strong: S-to-S and Weak-to-Weak: W-to-W) in order to account for GC-biased gene conversion (gBGC) (Bolívar et al., 2018).

Estimates of *d*_N_/*d*_S_ were obtained for collared flycatcher based on one-to-one orthologues between collared flycatcher and zebra finch (*Taeniopygia guttata*), using chicken (*Gallus gallus*) sequences as an outgroup. For this purpose, one-to-one orthologues were downloaded from Ensembl version 104. Based on the collared flycatcher reference genome, we identified 7559 genes located on autosomes and 303 genes on the Z-chromosome. For each gene, we then aligned the orthologous sequences across the three species using PRANK v170427 (Löytynoja, 2014) with help of a guide tree estimated with ClustalW v2.1 (Larkin et al., 2007). To estimate *d*_N_/*d*_S_, we used the software Bio++ v3.0 (Dutheil & Boussau, 2008), which first estimates a gene tree based on Maximum likelihood, and then maps substitutions based on stochastic mapping. The gene tree for each gene was estimated using a strand-symmetric L95 model (Lobry, 1995). After mapping substitutions, we separated substitutions into different categories, to estimate *d*_N_ and *d*_S_ using only GC-conservative changes in order to account for gBGC. To average *d*_N_ and *d*_S_ across genes, we weighted the counts for S-to-S and W-to-W substitutions by the proportion of GCs and ATs and took the sum of the two GC-conservative substitution types for nonsynonymous and synonymous substitutions.

We estimated the rate of adaptive substitutions, ω_a_, with the software DFE-alpha (Eyre-Walker & Keightley, 2009; Keightley & Eyre-Walker, 2007) using the divergence estimates described above and polymorphism data from all four species separately. Using GC-conservative sites, we obtained the site frequency spectrum (SFS) for 0-fold and 4-fold degenerate sites for each species. We then estimated the distribution of fitness effects (DFE) based on the 2-epoch model of population size change implemented in DFE-alpha. This resulted in one ω_a_ estimate based on polymorphisms within each species.

We obtained confidence intervals for *d*_N_/*d*_S_, *d*_N_/*d*_S_, and ω_a_ by randomly resampling genes with replacement and re-estimating each statistic for 100 bootstrap replicates.

### Population genomics statistics

We estimated *F*_*ST*_ for the two sister species pairs, collared and pied flycatcher and red-breasted and taiga flycatcher, for 200-kb genomic windows along the Z-chromosome. Following (Chase & Mugal, 2022), *F*_*ST*_ was estimated using VCFtools (Danecek et al., 2011). We identified *F*_*ST*_ peaks by Z transforming *F*_*ST*_ values for each chromosome, and applying a Savitzky-Golay filter to the transformed values. Windows with a smoothed Z-*F*_*ST*_ value above two were then identified as an *F*_*ST*_ peak. Estimates for *F*_*ST*_ along the autosomes were retrieved from (Chase & Mugal, 2022).

In addition to estimating *F*_*ST*_, we performed a selective sweep scan along the Z-chromosome to look for signatures of positive selection. We used the program SweepFinder2 (DeGiorgio et al., 2016) to implement the composite likelihood ratio (CLR) test (Nielsen et al., 2005), using the polarized SNP data for all four species individually. SweepFinder2 was run using the -ug option with a pre-computed background SFS for the Z-chromosome for each species and a user-defined grid with the location for each variant. Sites were first filtered to remove positions fixed for the ancestral allele within each species. We determined the significance threshold for the CLR test based on simulations in SLiM 3 (Haller & Messer, 2019). We simulated background selection occurring across an approximately 21 Mbp chromosome, based on gene density and recombination rate estimates from the collared flycatcher. The significance threshold based on these simulations is 46.25. We merged adjacent sites with significant CLR values into a single sweep region, and removed sweeps that contained only one position, or that had a site density less than 1bp/1kb, and then obtained presence/absence of selective sweeps in 200-kb windows. Estimates for selective sweeps along the autosomes were retrieved from (Chase & Mugal, 2022).

### Identification of fixed differences and shared polymorphisms

We identified sites that showed fixed differences between collared and pied flycatcher and between red-breasted and taiga flycatcher, as well as sites that displayed shared polymorphisms between the two species comparisons. We overlapped fixed differences with different functional categories, and compared the relative proportions of fixed differences in each category on the Z-chromosome compared to the autosomes. Functional categories included intergenic regions, intronic regions, conserved non-coding elements (CNEs) (Craig et al., 2018), untranslated regions (UTRs), four-fold degenerate sites, and zero-fold degenerate sites, where the latter three are based on the collared flycatcher annotation (Ensembl v. 104). Additionally, we examined whether any nonsynonymous fixed differences overlapped with a signature of selective sweeps in either of the two species compared.

### Estimates of gene flow

Collared and pied flycatcher have partially overlapping breeding ranges, and produce hybrid offspring in those contact zones. F1 hybrids are generally found to be sterile (Ålund et al., 2013; Svedin et al., 2008). Nevertheless, previous demographic modeling suggests a recent history of gene flow between the two species (Nadachowska-Brzyska et al., 2013; Nater et al., 2015). Less is known about the red-breasted and the taiga flycatcher (Hung & Zink, 2014; Svensson et al., 2005). For this reason and due to limited data availability of relevant reference species for the red-breasted and the taiga flycatcher, we here focus on gene flow between the collared and pied flycatcher. Specifically, we compared rates of gene flow between collared and pied flycatcher on the autosomes compared to the Z-chromosome using Patterson’s D statistic (Green et al., 2010). This test takes a four-taxon comparison, (P1,P2,P3,O), where O represents an outgroup, and can test for gene flow between population 1 (P1) and population 3 (P3) or between population 2 (P2) and P3. We estimated Patterson’s D for multiple population comparisons, which allowed us to investigate at what stage during the divergence of collared and pied flycatchers gene flow occurred. First, we set Atlas flycatcher as P1 and set Öland pied flycatcher and collared flycatcher as P2 and P3 respectively, and performed a separate test with Spanish pied flycatcher and Italian collared flycatcher as P2 and P3. Second, we used Spanish pied flycatcher as P1, Öland pied flycatcher as P2 and Öland collared flycatcher as P3, to determine whether there was greater evidence for gene flow between Öland populations. Third, we performed the test for Italian collared flycatcher as P1, Öland collared flycatcher as P2, and Öland pied flycatcher as P3. For each comparison, we used red-breasted flycatcher and snowy-browed flycatcher as outgroups. Polarized polymorphism data for all four species were obtained from previously published work (Burri et al., 2015).

We estimated Patterson’s D for autosomes and the Z-chromosome separately, using the derived allele frequencies (*p*) in the three in-group species with the formula:

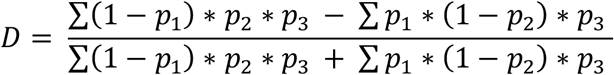

To test whether the estimates of Patterson’s D calculated were significantly different from zero, and thus showing a signature of gene flow, we performed jackknife resampling, removing blocks of 200-kb to estimate standard error and a Z-score for both the autosomes and the Z-chromosome.

In addition to autosomal and Z-chromosome average estimates of gene flow, we performed a window-based analysis to obtain local estimates of gene flow across the genome. We estimated the *f*_*d*_ statistic (Martin et al., 2015), which was developed to account for the high variance observed in the D-statistic in small genomic regions. We estimated this statistic only for the species comparisons that showed statistically significant estimates of gene flow on both autosomes and the Z-chromosome, since the *f*_*d*_ statistic is designed to detect gene flow between P2 and P3 and is not biologically meaningful when negative. We then identified genomic windows showing significantly reduced gene flow by first applying a smoothing algorithm to the window-based estimates of *f*_*d*_ for each chromosome, and then Z-transforming the smoothed estimates using the genome-wide mean and standard deviation. Windows with z-scores of -2 or lower were identified as significant outliers.

### Statistical analysis

All statistical analysis was performed in R version 4.0.3 (R Core Team 2020).

## Results

### Z/A ratio of effective population size in four *Ficedula* flycatcher species

As an estimate of *N*_e_ in the Z-chromosome compared to the autosomes, we calculated genome-wide π_Z_/π_A_ (All sites), π_Z_/π_A_ based on GC-conservative sites (GC cons), π_Z_/π_A_ after masking sites potentially affected by linked selection (No LS), as well as the ratio of historical *N*_e_ over 1Mya for all four species and then corrected those ratios for differences in mutation rate among the Z-chromosome and autosomes (Irwin, 2018) (Fig. 1A; Tables S2 & S3). Estimates of the Z:A ratio in *N*_e_ were largely consistent for π_Z_/π_A_ (All sites), π_Z_/π_A_ (GC cons), π_Z_/π_A_ (No LS) and PSMC_Z/A_. Collared flycatcher and pied flycatcher showed a ratio clearly < 0.75, while red-breasted flycatcher and taiga flycatcher showed larger values close to 0.75. The consistency among π_Z_/π_A_ estimates suggests that differences in the strength of gBGC and/or linked selection between the Z-chromosome and the autosomes do not show an impact on the Z:A ratio of *N*_e_ in *Ficedula* flycatchers. To investigate if the differences in the Z:A ratio among species are associated with differences in recent demographic histories, we visualize their PSMC historical population size (Fig. 1B). This showed no clear association. Recent demographic histories were more similar among collared flycatcher and red-breasted flycatcher and among pied flycatcher and taiga flycatcher, than among collared flycatcher and pied flycatcher and among red-breasted flycatcher and taiga flycatcher. Variance of reproductive success in males, on the other hand, reflects the observed differences in the Z:A ratio among species. While collared flycatchers and pied flycatchers are partly polygynous, red-breasted flycatchers (and taiga flycatchers) are purely monogamous (Storchová & Hořák, 2018).

**Figure 1:**
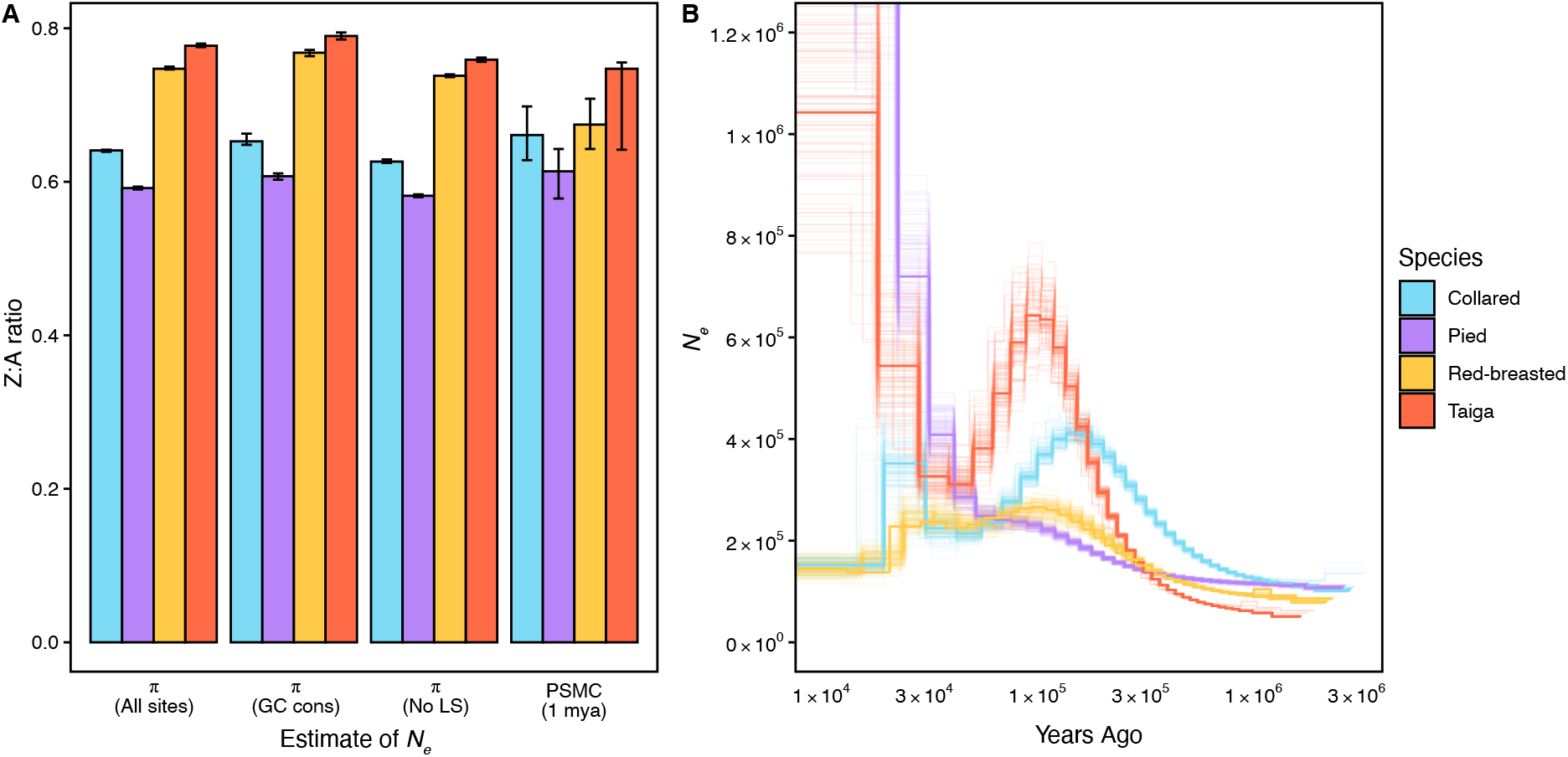
Effective population size (*N*_e_) on the autosomes and the Z-chromosome. Panel **A** shows the ratios of different estimates of *N*_e_ for all four species on the Z-chromosome versus the autosomes (Z:A). Estimates of *N*_e_ are based on nucleotide diversity (π) using all SNPs (All sites), π corrected for linked selection by masking protein-coding sequences and conserved non-coding elements (CNEs) and their 1kb flanking regions (No LS), and estimates based on PSMC historical population size for the last 1Mya. All estimates are corrected for male-biased mutation rate. See Tables S2 and S3 for respective estimates for the Z-chromosome and autosomes. Panel **B** shows historical changes in population size for all four species estimated on the autosomes. The bold line represents the genome-wide estimate, bootstrap replicates are shown in lighter colour. See Fig S1 for estimates on the Z-chromosome.

### No evidence that adaptation drives the *fast-Z* effect in *Ficedula* flycatchers

Fig 1A illustrates that *N*_e_ is smaller on the Z-chromosome than the autosomes in all four species. We therefore examined if these differences in *N*_e_, and hence the strength of genetic drift, and/or the hemizygote state of the Z-chromosome in females influence the efficacy of natural selection. For this purpose, we estimated several measures of the strength of selection on protein-coding sequences on the Z-chromosome compared to the autosomes. We observed that *d*_N_/*d*_S_ was higher on the Z-chromosome compared to the autosomes for all four species (Table 1). Similarly, branch-specific *d*_N_/*d*_S_ estimates for the flycatcher lineage after the split from zebra finch was higher on the Z-chromosome, which is consistent with a *fast-Z* effect in *Ficedula* flycatchers. To assess if elevated *d*_N_/*d*_S_ is a result of relaxed purifying selection or stronger positive selection on the Z-chromosome, we estimated the adaptive substitution rate ω_a_. We used polymorphism data of each of the four species separately to estimate the distribution of fitness effects, which provides information on the influence of demography on ω_a_ estimates. This revealed that estimates of ω_a_ were lower on the Z-chromosome compared to the autosomes for pied flycatcher and taiga flycatcher (Table 1), which both show a recent increase in population size (Fig. 1B). Collared flycatcher and red-breasted flycatcher, on the other hand, which both show less fluctuations in recent population size (Fig. 1B), showed higher or not significantly different estimates on the Z-chromosome compared to the autosomes. These differences in ω_a_ estimates among species are solely governed by differences in the SFS among species, which is also apparent in differences in *d*_N_/*d*_S_ estimates. The impact of demography on estimates of ω_a_ therefore makes it difficult to assess the role of adaptation in the *fast-Z* effect.

**Table 1:** Estimates of selection on coding sequences for autosomes and the Z-chromosome. Shown are estimates of *d*_N_/*d*_S_, *d*_N_/*d*_S_, and ω_a_ separately for genes on the autosomes and on the Z-chromosome. All estimates are based on GC-conservative sites only. See Table S4 for estimates based on all sites.

|  | Collared | Pied | Red-breasted | Taiga |
| --- | --- | --- | --- | --- |
| $\pi_N/\pi_S$ A | 0.174<br>[0.159;0.186] | 0.186<br>[0.166;0.209] | 0.164<br>[0.148;0.175] | 0.169<br>[0.155;0.181] |
| $\pi_N/\pi_S$ Z | 0.183<br>[0.116;0.262] | 0.344<br>[0.163;0.653] | 0.194<br>[0.139;0.284] | 0.229<br>[0.166;0.309] |
| $d_N/d_S$ A | 0.171 [0.164;0.177] | | | |
| $d_N/d_S$ Z | 0.198 [0.166;0.233] | | | |
| $\omega_a$ A | 0.0592<br>[0.0434;0.0767] | 0.0422<br>[0.0199;0.0668] | 0.0451<br>[0.0259;0.0622] | 0.0900<br>[0.0771;0.100] |
| $\omega_a$ Z | 0.112<br>[0.0509;0.187] | 0.0182<br>[-0.0896;0.107] | 0.0474<br>[-0.0533;0.129] | 0.0594<br>[-0.0274;0.104] |

To complement the analysis based on estimates of selection on protein-coding sequences, we estimated window-based *F*_*ST*_ on the Z-chromosome in 200-kb windows for the two sister species comparisons: collared and pied flycatchers and red-breasted and taiga flycatchers (Fig 2A & 2D). For both comparisons *F*_*ST*_ was notably higher on the Z-chromosome than on the autosomes, with Z-chromosome average *F*_*ST*_ 0.55 vs. 0.29 for the autosomes between collared and pied flycatcher and 0.70 vs. 0.62 between red-breasted and taiga flycatcher. We identified *F*_*ST*_ peaks on both the autosomes and the Z-chromosome, and found that for red-breasted and taiga flycatcher, *F*_*ST*_ peaks occurred more frequently on autosomes (Table 2), while there was no significant difference for collared and pied flycatcher. Thus, the higher *F*_*ST*_ levels on the Z-chromosome appeared to be a chromosome-wide effect as a result of faster lineage sorting of rather than more prevalent signatures of *F*_*ST*_ peaks. Since selective sweeps have been found to play a role in shaping *F*_*ST*_ peaks in Ficedula flycatchers (Chase et al., 2021), the low prevalence of *F*_*ST*_ peaks on the Z-chromosome in fact suggests that linked positive selection is not more common on the Z-chromosome compared to the autosomes.

**Table 2:** Number of *F*_*ST*_ peaks on autosomes compared to the Z-chromosome. Shown are the number of 200-kb windows overlapping with an *F*_*ST*_ peak or not for both species comparisons, for both the autosomes and the Z-chromosome. Odds ratio and *P*-value indicate the significance level of the overrepresentation on the autosomes based on Fisher exact tests.

| | Autosome | Z-chromosome | Odds ratio | $P$-value |
| --- | --- | --- | --- | --- |
| **Coll/pied $F_{ST}$ peak** | 179 | 1 | 2.2 | 0.10 |
| **Coll/pied no $F_{ST}$ peak** | 4517 | 278 |  |  |
| **Red-breasted/taiga $F_{ST}$ peak** | 247 | 6 | 2.6 | 0.017 |
| **Red-breasted/taiga no $F_{ST}$ peak** | 4449 | 277 |  |  |

**Figure 2:**
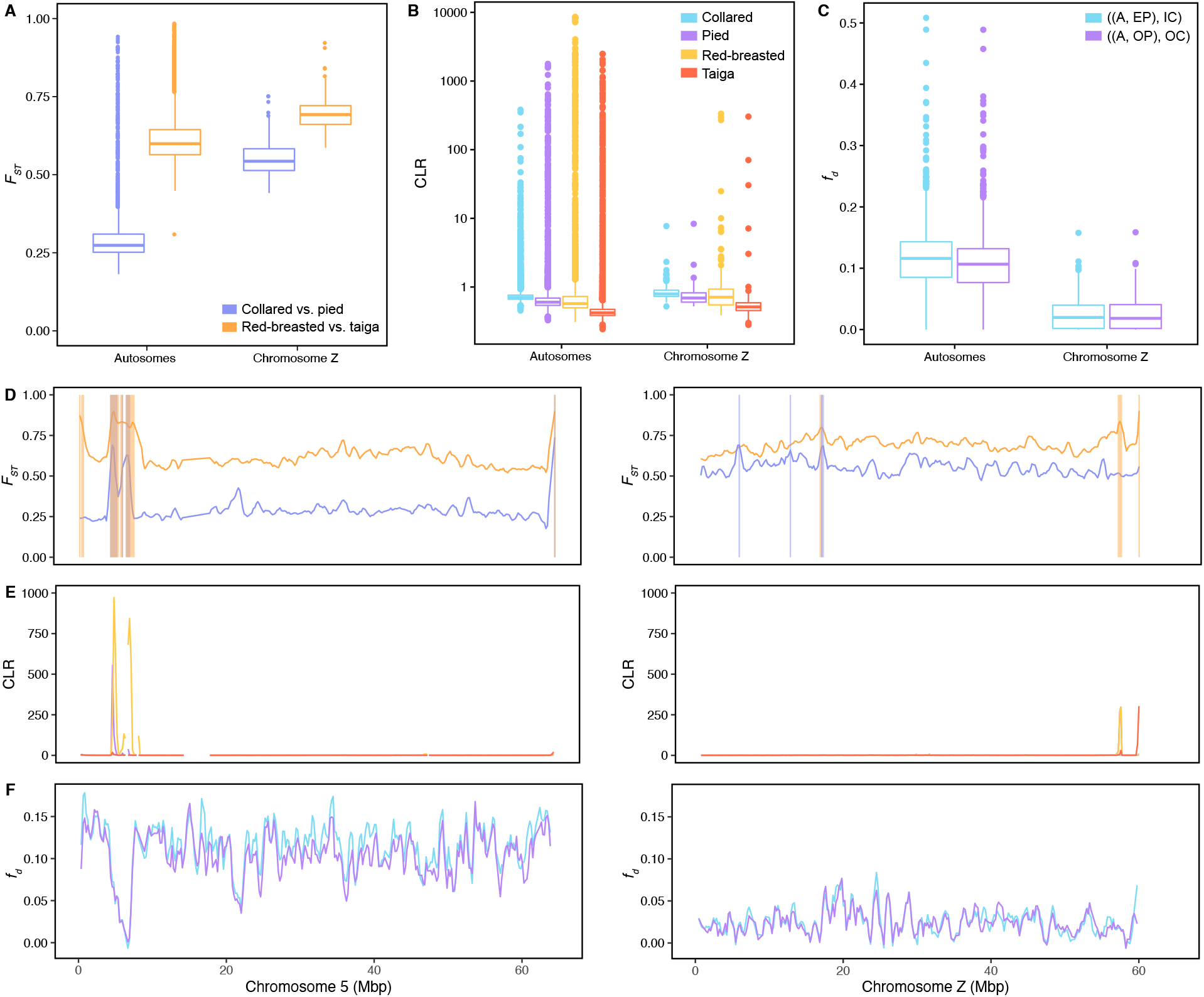
Signatures of linked selection and gene flow on the Z-chromosome compared to autosomes. Panel **A** shows the distributions of window-based estimates of *F*_ST_ between collared and pied flycatchers (purple) and red-breasted and taiga flycatchers (orange) separately for autosomes and the Z-chromosome. Panel **B** shows window-based CLR estimates for the four species in log-scale. Panel **C** shows window-based estimates of *f*_*d*_ for gene flow estimates between Spanish pied flycatcher and Italian collared flycatcher (blue) and between Öland pied flycatcher and Öland collared flycatcher (purple) using atlas flycatcher as a reference. Panel **D** shows measures of *F*_*ST*_ estimated in 200-kb genomic windows for chromosome 5 (left) and the Z-chromosome (right). Shaded rectangles demonstrate the locations of significant *F*_*ST*_ peaks in both species comparisons. Panel **E** shows CLR estimates for collared (blue), pied (purple), red-breasted (yellow) and taiga (red) flycatcher on both chromosomes. Panel **F** shows window-based estimates of *f*_*d*_ for gene flow estimates between Spanish pied flycatcher and Italian collared flycatcher (blue) and between Öland pied flycatcher and Öland collared flycatcher (purple). See Figure S2 for estimates of *F*_ST_ for all chromosomes, Figure S3 for estimates of CLR for all chromosomes, and Figure S4 for estimates of *f*_*d*_ along all chromosomes.

In order to corroborate this finding, we also compared selective sweep scans in each species on the autosomes and Z-chromosome (Fig 2B & 2E). Consistent with observations for *F*_*ST*_ peaks, this revealed that red-breasted and taiga flycatchers showed a greater proportion of selective sweeps on the autosomes than the Z-chromosome, while collared flycatcher and pied flycatchers showed no significant difference between the autosomes and the Z-chromosome (Table 3). In addition, we found that selective sweep signatures on the Z-chromosome overlapped on average with a lower fraction of nonsynonymous fixed differences compared to the autosomes (Table S5).

**Table 3:** Prevalence of selective sweeps on autosomes compared to the Z-chromosome. Shown are the number of 200-kb windows overlapping with a significant selective sweep signature on the autosomes and the Z-chromosome compared to the number of 200-kb windows not overlapping with a selective sweep. Odds ratio and *P*-value indicate the significance level of the overrepresentation on the autosomes based on Fisher exact tests.

|  | **Autosome** | **Z-chromosome** | **Odds ratio** | ***P*-value** |
| --- | --- | --- | --- | --- |
| **Collared sweep** | 59 | 1 | 4.2 | 0.18 |
| **Collared no sweep** | 4715 | 335 |  |  |
| **Pied sweep** | 137 | 6 | 1.6 | 0.30 |
| **Pied no sweep** | 4588 | 327 |  |  |
| **Red-breasted sweep** | 248 | 9 | 2.0 | 0.039 |
| **Red-breasted no sweep** | 4512 | 332 |  |  |
| **Taiga sweep** | 223 | 5 | 3.3 | 0.0027 |
| **Taiga no sweep** | 4623 | 341 |  |  |

We next computed the density of fixed differences between both collared and pied flycatcher and red-breasted and taiga flycatcher in six different functional regions of the genome, i.e., intergenic and intronic regions, UTRs, CNEs, as well as four-fold and zero-fold degenerate sites. We then compared the functional overlap of the fixed differences on the autosomes and the Z-chromosome. For both species pairs, we observed the highest density of fixed differences in introns and intergenic regions on both the autosomes and the Z-chromosome, followed by four-fold degenerate sites and UTRs (Fig 3). Both CNEs and zero-fold degenerate sites showed the lowest density of fixed differences for both species pairs and chromosome types (Fig 3). Overall, there was no observable increase of fixed differences in functional categories potentially evolving under selective constraint (UTRs, CNEs, and zero-fold degenerate sites) versus potentially neutrally evolving categories (intronic regions, intergenic regions, and four-fold degenerate sites) for the Z-chromosome versus autosomes (Table S6). These results suggest that the increase in fixed differences on the Z-chromosome is driven primarily by faster lineage-sorting rather than positive selection. In line with this conclusion, we also find fewer shared polymorphisms between the two species pairs on the Z-chromosome compared to autosomes (Table S7).

**Figure 3:**
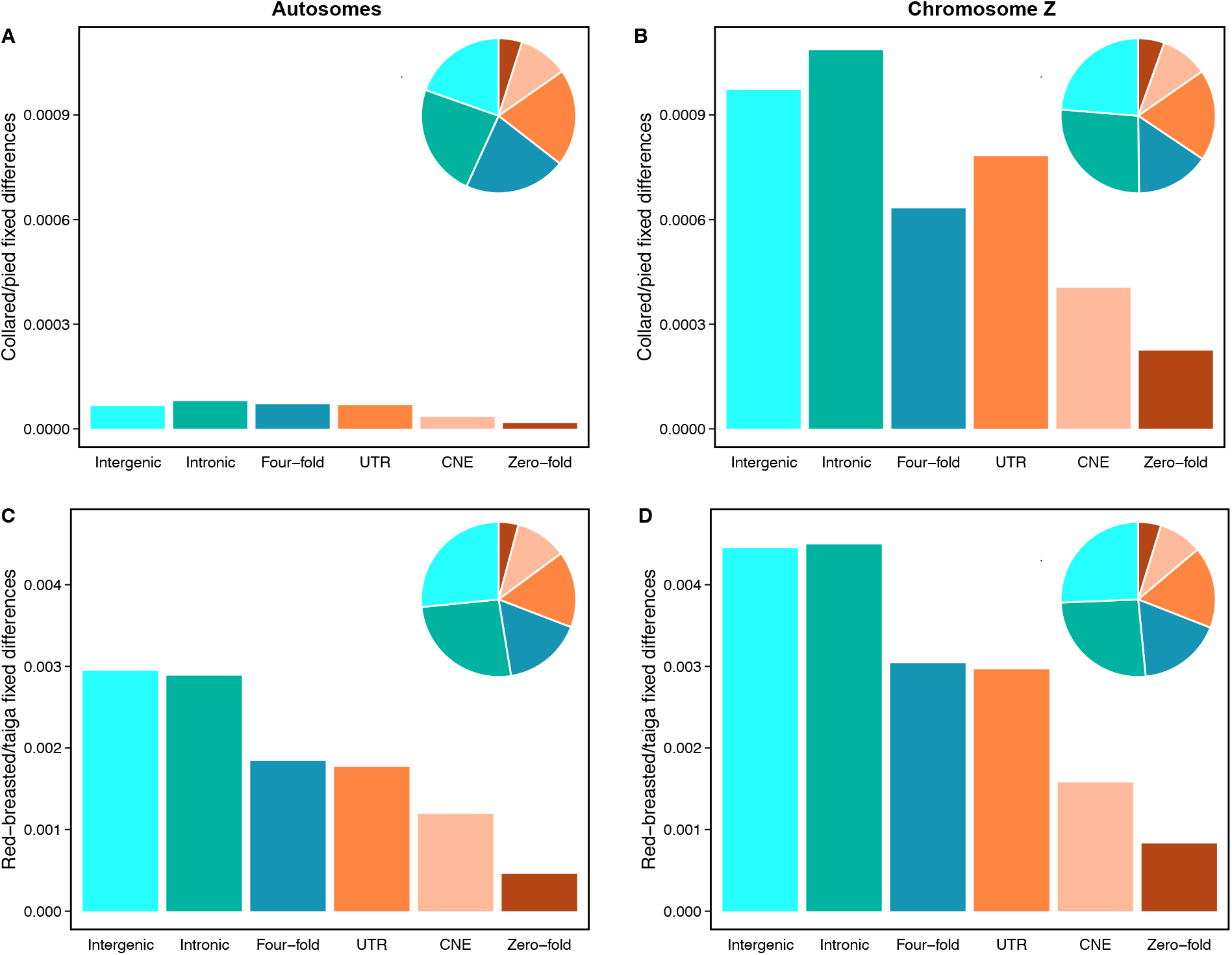
Functional overlap of fixed differences on the Z-chromosome versus the autosomes. Panels **A** and **B** show the densities of fixed differences in different functional regions between collared and pied flycatcher on the autosomes and the Z-chromosome, respectively. Panels **C** and **D** show the same between red-breasted and taiga flycatcher. In all four panels, fixed differences are grouped into the following functional categories: intergenic, intronic, and untranslated regions (UTRs), conserved noncoding elements (CNEs), four-fold degenerate sites, and zero-fold degenerate sites. Colors correspond to whether differences are suggested to be evolving under selective constraint (orange shade) or neutrally (blue shade). The pie chart insets show the proportion of fixed differences in each functional category.

### Reduced signatures of introgression on the Z-chromosome

We performed ABBA-BABA tests to detect signatures of introgression for several combinations of populations of collared flycatcher and pied flycatcher with different reference species. First, we tested for gene flow between the pair of Öland collared flycatcher and Öland pied flycatcher as well as the pair of Italian collared flycatcher and Spanish pied flycatcher, using Atlas flycatcher as a reference species. This revealed that in both settings there was a significant excess of ABBA sites compared to BABA sites, which provides evidence for a history of gene flow between collared flycatcher and pied flycatcher (Table 4). The Z-chromosome showed a lower value of the D-statistic compared to the autosomes (Table 4), suggesting there has been a relative reduction in gene flow on the Z-chromosome. We also tested for evidence of gene flow between the Öland populations of collared flycatcher and pied flycatcher using either Italian collared flycatcher or Spanish pied flycatcher as reference species. For both settings, there was no significant evidence of gene flow on the Z-chromosome, and a significant but small effect size of gene flow on the autosomes (Table 4). Taken together, these D-statistic estimates point to a more ancient history of gene flow between collared flycatcher and pied flycatcher, which occurred before the divergence of the different collared flycatcher and pied flycatcher populations considered here, and that gene flow is lower on the Z-chromosome compared to the autosomes.

**Table 4:** Signatures of gene flow on the autosomes compared to the Z-chromosome. Shown are results from ABBA-BABA tests on the autosomes and Z-chromosome for five different population/species comparisons. The populations/species are listed in the format ((P1, P2), P3), where P1 and P2 represent sister populations/species, and where we are testing for evidence of gene flow between either P1 or P2 with P3. Populations/species included in the tests are Atlas flycatcher (A), Öland pied flycatcher (OP), Spanish pied flycatcher (EP), Öland collared flycatcher (OC), and Italian collared flycatcher (IC). *P*-values were estimated by block jackknife resampling, removing 200-kb windows.

| Species | Chromosome | D-statistic (±se) | Z-score | *P*-value |
| --- | --- | --- | --- | --- |
| ((A, OP), OC) | Autosomes | 0.18 (± 0.0012) | 145 | $< 10^{-12}$ |
| ((A, OP), OC) | Chromosome Z | 0.10 (± 0.0089) | 11 | $< 10^{-12}$ |
| ((A, EP), IC) | Autosomes | 0.18 (± 0.0012) | 145 | $< 10^{-12}$ |
| ((A, EP), IC) | Chromosome Z | 0.099 (± 0.0086) | 12 | $< 10^{-12}$ |
| ((EP, OP), OC) | Autosomes | 0.0018 (± 0.00042) | 4.3 | $1.7 \cdot 10^{-5}$ |
| ((EP, OP), OC) | Chromosome Z | -0.0022 (± 0.0060) | -0.36 | 0.72 |
| ((IC, OC), OP) | Autosomes | -0.0015 (± 0.00040) | -3.7 | $2.2 \cdot 10^{-4}$ |
| ((IC, OC), OP) | Chromosome Z | 0.00027 (± 0.0037) | 0.072 | 0.94 |

We next estimated window-based signatures of gene flow using the *f*_*d*_ statistic for the two comparisons that showed signatures of gene flow on both autosomes and the Z-chromosome. Consistent with genome-wide ABBA-BABA tests, this revealed that the distribution of *f*_*d*_ values on the Z-chromosome was lower than on the autosomes (Fig 2). This observation was apparent for all autosomal windows combined, and for windows separately for each chromosome, with the exception of some microchromosomes with few data points (Fig S4). The same trend was observed whether we compared Öland collared flycatcher and pied flycatcher populations or Italian collared flycatcher and Spanish pied flycatcher populations. We identified many windows on the Z-chromosome showing significantly reduced estimates of *f*_*d*_ compared to the genome-wide average, and we found that the Z-chromosome was enriched for these windows (Table S8).

Since the Z-chromosome has on average lower rates of recombination compared to autosomes (Kawakami et al., 2014), the reduction in recombination rate alone could potentially explain the reduced signatures of gene flow we detect on the Z-chromosome (Martin et al., 2019; Schumer et al., 2018). We therefore examined the relationship between recombination rate and signatures of introgression by performing a linear regression on pedigree-based recombination rate estimated in collared flycatcher (Kawakami et al., 2014) and the *f*_*d*_ statistic for both population comparisons. Although there was a significant relationship for both comparisons (Öland collared flycatcher and pied flycatcher: R^2^ = 0.020, *P*-value = 1.6∙10^−13^; Italian collared flycatcher and Spanish pied flycatcher: R^2^ = 0.019, *P*-value = 9.0∙10^−13^), the low R^2^ demonstrates that recombination rate explains little of the genome-wide variation in *f*_*d*_.

## Discussion

Our analysis of the Z:A ratio of effective population size (*N*_e_) reveals that collared flycatcher and pied flycatcher show a relatively lower *N*_e_ on the Z-chromosome than red-breasted flycatcher and taiga flycatcher. The Z:A ratio of *N*_e_ was significantly lower than 0.75 in the two black-and-white flycatchers, while red-breasted flycatcher and taiga flycatcher showed a ratio close to the expectation of 0.75 for equal sex ratios. These differences relate very well to the mating behaviors reported for different *Ficedula* flycatchers (Storchová & Hořák, 2018). Collared flycatchers and pied flycatchers are partly polygynous, while red-breasted flycatchers (and taiga flycatchers) are purely monogamous, where a greater reproductive variance in males is expected to reduce the Z:A ratio of *N*_e_ (Vicoso & Charlesworth, 2009). Differences in the strength of linked selection among the Z-chromosome and autosomes, on the other hand, appear not to show any strong influence on the Z:A ratio of *N*_e_ in the *Ficedula* flycatcher lineage. The lack of a strong impact of linked selection on the Z:A ratio of *N*_e_ suggests caution in the presumption that linked selection might explain low values of Z:A diversity observed in birds (Irwin, 2018), which is further supported by our observation that selective sweep signatures are not more pronounced on the Z-chromosome. Also, differences in the demographic history between the four species did not correlate with observed differences in the Z:A ratio of *N*_e_ among species. It therefore appears that life-history traits and mating behavior are the strongest predictors of differences in the Z:A ratio of *N*_e_ among *Ficedula* flycatchers.

Despite the observed differences in the Z:A ratio of *N*_e_ among species, *N*_e_ is clearly smaller on the Z-chromosome than the autosomes for all four *Ficedula* flycatchers. The stronger impact of genetic drift on the Z-chromosome than the autosomes therefore needs to be considered in the evaluation of the driving forces of the *fast-Z* and *large-Z* effects. Indeed, macro- and micro-evolutionary signatures of natural selection suggest that genetic drift rather than adaptation explains the *fast-Z* effect in *Ficedula* flycatchers, which is in line with previous observations in birds (Hayes et al., 2020; Mank et al., 2010; Wang et al., 2014). We find evidence for reduced purifying selection on the Z-chromosome, but no evidence for a stronger signature of positive selection on the Z-chromosome than the autosomes. On the contrary, if anything, the signature of positive selection appears to be weaker on the Z-chromosome than the autosomes. While divergent demographic history results in inconsistent patterns of the rate of adaptive evolution between the Z-chromosome and the autosomes among species, comparison of the prevalence of selective sweep signatures provides a clearer picture. For collared flycatcher and pied flycatcher no significant difference in the prevalence of selective sweeps could be found between the Z-chromosome and the autosomes. For red-breasted flycatcher and taiga flycatcher, on the other hand, selective sweep signatures were clearly less prevalent on the Z-chromosome than the autosomes.

Yet, despite the lack of stronger positive selection on the Z-chromosome, we observe evidence for reduced gene flow on the Z-chromosome compared to the autosomes. Thus, our results support the hypothesis that the *large-Z* effect does not necessarily need to invoke positive or divergent selection. The reduction in gene flow appears to be a chromosome-wide effect rather than limited to narrow barrier loci, which is in good agreement with the chromosome-wide effect of genetic drift. This mechanism can further explain the presence of genomic signatures of a *large-Z* effect despite of a lack of ‘active’ differential introgression on the Z-chromosome (Hogner et al., 2012). Specifically, the relative reduction in *N*_e_ on the Z-chromosome compared to the autosomes leads to faster lineage sorting and elevated *F*_ST_ on the entire Z-chromosome, where the latter is frequently perceived as evidence for *fast-Z* and *large-Z* effects. However, even though we don’t find signatures of adaptation on the Z-chromosome, our analysis does not exclude the possibility that Z-linked loci could play an important role for hybrid incompatibilities and reproductive isolation. The accelerated differentiation of the Z-chromosome could potentially lead to an accelerated accumulation of incompatibilities between the Z-chromosome and interacting loci on the autosomes. It is in fact tempting to speculate that such a mechanism could trigger a snowball effect of selective sweeps on the autosomes, which could be the driving mechanism of hybrid sterility between collared and pied flycatcher hybrids (Segami et al., 2022).

## Acknowledgements

The authors thank Laure Ségurel for feedback on an earlier version of the manuscript. The computational infrastructure and data storage were enabled by resources provided by the Swedish National Infrastructure for Computing (SNIC), partially funded by the Swedish Research Council through grant agreement No. 2018-05973. MAC and CFM have conducted their work with financial support from the Knut and Alice Wallenberg Foundation (2014/0044 to Hans Ellegren) and the Swedish Research Council (2013-8271 to Hans Ellegren).

## Author contributions

MAC and CFM conceived the study. MAC conducted data analyses. MV contributed to the data analyses. CFM supervised the study. MAC and CFM wrote the manuscript. All authors critically read and approved the final version of the manuscript.

## Data accessibility

Sequencing data for all samples are available at the EMBL-EBI European Nucleotide Archive (ENA; http://www.ebi.ac.uk/ena) with the following accession numbers: PRJEB43825 (taiga and red-breasted flycatchers), PRJEB22864 (collared flycatcher), and PRJEB7359 (pied and snowy-browed flycatchers). VCF files used for analyses will be deposited on dryad upon publication. Scripts used for analysis will be publicly available on GitHub upon publication.

## References

Ålund, M., Immler, S., Rice, A. M., & Qvarnström, A. (2013). Low fertility of wild hybrid male flycatchers despite recent divergence. Biology Letters, 9(3), 20130169. https://doi.org/10.1098/rsbl.2013.0169

Avery, P. J. (1984). The population genetics of haplo-diploids and X-linked genes. Genetics Research, 44(3), 321–341. https://doi.org/10.1017/S0016672300026550

Axelsson, E., Smith, N. G. C., Sundström, H., Berlin, S., & Ellegren, H. (2004). Male-Biased Mutation Rate and Divergence in Autosomal, Z-Linked and W-Linked Introns of Chicken and Turkey. Molecular Biology and Evolution, 21(8), 1538–1547. https://doi.org/10.1093/molbev/msh157

Bolívar, P., Mugal, C. F., Rossi, M., Nater, A., Wang, M., Dutoit, L., & Ellegren, H. (2018). Biased Inference of Selection Due to GC-Biased Gene Conversion and the Rate of Protein Evolution in Flycatchers When Accounting for It. Molecular Biology and Evolution, 35(10), 2475–2486. https://doi.org/10.1093/molbev/msy149

Borge, T., Webster, M. T., Andersson, G., & Saetre, G.-P. (2005). Contrasting Patterns of Polymorphism and Divergence on the Z Chromosome and Autosomes in Two Ficedula Flycatcher Species. Genetics, 171(4), 1861–1873. https://doi.org/10.1534/genetics.105.045120

Burri, R., Nater, A., Kawakami, T., Mugal, C. F., Olason, P. I., Smeds, L., Suh, A., Dutoit, L., Bureš, S., Garamszegi, L. Z., Hogner, S., Moreno, J., Qvarnström, A., Ružić, M., Sæther, S.-A., Sætre, G.-P., Török, J., & Ellegren, H. (2015). Linked selection and recombination rate variation drive the evolution of the genomic landscape of differentiation across the speciation continuum of Ficedula flycatchers. Genome Research, 25(11), 1656–1665. https://doi.org/10.1101/gr.196485.115

Charlesworth, B., Coyne, J. A., & Barton, N. H. (1987). The Relative Rates of Evolution of Sex Chromosomes and Autosomes. The American Naturalist, 130(1), 113–146. https://doi.org/10.1086/284701

Chase, M. A., Ellegren, H., & Mugal, C. F. (2021). Positive selection plays a major role in shaping signatures of differentiation across the genomic landscape of two independent Ficedula flycatcher species pairs*. Evolution, 75(9), 2179–2196. https://doi.org/10.1111/evo.14234

Chase, M. A., & Mugal, C. F. (2022). The role of recombination dynamics in shaping signatures of direct and indirect selection across the Ficedula flycatcher genome(p. 2022.08.11.503468). bioRxiv. https://doi.org/10.1101/2022.08.11.503468

Coyne, J. A. (1984). Genetic basis of male sterility in hybrids between two closely related species of Drosophila. Proceedings of the National Academy of Sciences, 81(14), 4444–4447. https://doi.org/10.1073/pnas.81.14.4444

Coyne, J. A. (2018). “Two Rules of Speciation” revisited. Molecular Ecology, 27(19), 3749– 3752. https://doi.org/10.1111/mec.14790

Craig, R. J., Suh, A., Wang, M., & Ellegren, H. (2018). Natural selection beyond genes: Identification and analyses of evolutionarily conserved elements in the genome of the collared flycatcher (Ficedula albicollis). Molecular Ecology, 27(2), 476–492. https://doi.org/10.1111/mec.14462

Danecek, P., Auton, A., Abecasis, G., Albers, C. A., Banks, E., DePristo, M. A., Handsaker, R. E., Lunter, G., Marth, G. T., Sherry, S. T., McVean, G., Durbin, R., & Group, 1000 Genomes Project Analysis. (2011). The variant call format and VCFtools. Bioinformatics, 27(15), 2156–2158. https://doi.org/10.1093/bioinformatics/btr330

DeGiorgio, M., Huber, C. D., Hubisz, M. J., Hellmann, I., & Nielsen, R. (2016). S weep F inder 2: Increased sensitivity, robustness and flexibility. Bioinformatics, 32(12), 1895– 1897. https://doi.org/10.1093/bioinformatics/btw051

Dutheil, J., & Boussau, B. (2008). Non-homogeneous models of sequence evolution in the Bio++ suite of libraries and programs. BMC Evolutionary Biology, 8(1), 255. https://doi.org/10.1186/1471-2148-8-255

Ellegren, H. (2007). Characteristics, causes and evolutionary consequences of male-biased mutation. Proceedings of the Royal Society B: Biological Sciences, 274(1606), 1–10. https://doi.org/10.1098/rspb.2006.3720

Eyre-Walker, A., & Keightley, P. D. (2009). Estimating the Rate of Adaptive Molecular Evolution in the Presence of Slightly Deleterious Mutations and Population Size Change. Molecular Biology and Evolution, 26(9), 2097–2108. https://doi.org/10.1093/molbev/msp119

Feder, J. L., Egan, S. P., & Nosil, P. (2012). The genomics of speciation-with-gene-flow. Trends in Genetics, 28(7), 342–350. https://doi.org/10.1016/j.tig.2012.03.009

Green, R. E., Krause, J., Briggs, A. W., Maricic, T., Stenzel, U., Kircher, M., Patterson, N., Li, H., Zhai, W., Fritz, M. H.-Y., Hansen, N. F., Durand, E. Y., Malaspinas, A.-S., Jensen, J. D., Marques-Bonet, T., Alkan, C., Prüfer, K., Meyer, M., Burbano, H. A., … Pääbo, S. (2010). A Draft Sequence of the Neandertal Genome. Science, 328(5979), 710–722. https://doi.org/10.1126/science.1188021

Haller, B. C., & Messer, P. W. (2019). SLiM 3: Forward Genetic Simulations Beyond the Wright–Fisher Model. Molecular Biology and Evolution, 36(3), 632–637. https://doi.org/10.1093/molbev/msy228

Hayes, K., Barton, H. J., & Zeng, K. (2020). A Study of Faster-Z Evolution in the Great Tit (Parus major). Genome Biology and Evolution, 12(3), 210–222. https://doi.org/10.1093/gbe/evaa044

Hedrick, P. W. (2007). Sex: Differences in Mutation, Recombination, Selection, Gene Flow, and Genetic Drift. Evolution, 61(12), 2750–2771. https://doi.org/10.1111/j.1558-5646.2007.00250.x

Hogner, S., Sæther, S. A., Borge, T., Bruvik, T., Johnsen, A., & Sætre, G.-P. (2012). Increased divergence but reduced variation on the Z chromosome relative to autosomes in Ficedula flycatchers: Differential introgression or the faster-Z effect? Ecology and Evolution, 2(2), 379–396. https://doi.org/10.1002/ece3.92

Hung, C.-M., & Zink, R. M. (2014). Distinguishing the effects of selection from demographic history in the genetic variation of two sister passerines based on mitochondrial– nuclear comparison. Heredity, 113(1), Article 1. https://doi.org/10.1038/hdy.2014.9

Irwin, D. E. (2018). Sex chromosomes and speciation in birds and other ZW systems. Molecular Ecology, 27(19), 3831–3851. https://doi.org/10.1111/mec.14537

Kawakami, T., Smeds, L., Backström, N., Husby, A., Qvarnström, A., Mugal, C. F., Olason, P., & Ellegren, H. (2014). A high-density linkage map enables a second-generation collared flycatcher genome assembly and reveals the patterns of avian recombination rate variation and chromosomal evolution. Molecular Ecology, 23(16), 4035–4058. https://doi.org/10.1111/mec.12810

Keightley, P. D., & Eyre-Walker, A. (2007). Joint Inference of the Distribution of Fitness Effects of Deleterious Mutations and Population Demography Based on Nucleotide Polymorphism Frequencies. Genetics, 177(4), 2251–2261. https://doi.org/10.1534/genetics.107.080663

Kirkpatrick, M., & Hall, D. W. (2004). Male-Biased Mutation, Sex Linkage, and the Rate of Adaptive Evolution. Evolution, 58(2), 437–440. https://doi.org/10.1111/j.0014-3820.2004.tb01659.x

Larkin, M. A., Blackshields, G., Brown, N. P., Chenna, R., McGettigan, P. A., McWilliam, H., Valentin, F., Wallace, I. M., Wilm, A., Lopez, R., Thompson, J. D., Gibson, T. J., & Higgins, D. G. (2007). Clustal W and Clustal X version 2.0. Bioinformatics, 23(21), 2947–2948. https://doi.org/10.1093/bioinformatics/btm404

Leroy, T., Rousselle, M., Tilak, M.-K., Caizergues, A. E., Scornavacca, C., Recuerda, M., Fuchs, J., Illera, J. C., Swardt, D. H. D., Blanco, G., Thébaud, C., Milá, B., & Nabholz, B. (2021). Island songbirds as windows into evolution in small populations. Current Biology, 31(6), 1303–1310. https://doi.org/10.1016/j.cub.2020.12.040

Li, H., & Durbin, R. (2011). Inference of human population history from individual whole-genome sequences. Nature, 475(7357), 493–496. https://doi.org/10.1038/nature10231

Liu, S., & Hansen, M. M. (2017). PSMC (pairwise sequentially Markovian coalescent) analysis of RAD (restriction site associated DNA) sequencing data. Molecular Ecology Resources, 17(4), 631–641. https://doi.org/10.1111/1755-0998.12606

Lobry, J. R. (1995). Properties of a general model of DNA evolution under no-strand-bias conditions. Journal of Molecular Evolution, 40(3), 326–330. https://doi.org/10.1007/BF00163237

Löytynoja, A. (2014). Phylogeny-aware alignment with PRANK. In D. J. Russell (Ed.), Multiple Sequence Alignment Methods(pp. 155–170). Humana Press. https://doi.org/10.1007/978-1-62703-646-7_10

Mank, J. E., Nam, K., & Ellegren, H. (2010). Faster-Z Evolution Is Predominantly Due to Genetic Drift. Molecular Biology and Evolution, 27(3), 661–670. https://doi.org/10.1093/molbev/msp282

Martin, S. H., Davey, J. W., & Jiggins, C. D. (2015). Evaluating the Use of ABBA–BABA Statistics to Locate Introgressed Loci. Molecular Biology and Evolution, 32(1), 244– 257. https://doi.org/10.1093/molbev/msu269

Martin, S. H., Davey, J. W., Salazar, C., & Jiggins, C. D. (2019). Recombination rate variation shapes barriers to introgression across butterfly genomes. PLOS Biology, 17(2), e2006288. https://doi.org/10.1371/journal.pbio.2006288

Nachman, M. W., & Payseur, B. A. (2012). Recombination rate variation and speciation: Theoretical predictions and empirical results from rabbits and mice. Philosophical Transactions of the Royal Society B: Biological Sciences, 367(1587), 409–421. https://doi.org/10.1098/rstb.2011.0249

Nadachowska-Brzyska, K., Burri, R., & Ellegren, H. (2019). Footprints of adaptive evolution revealed by whole Z chromosomes haplotypes in flycatchers. Molecular Ecology, 28(9), 2290–2304. https://doi.org/10.1111/mec.15021

Nadachowska-Brzyska, K., Burri, R., Olason, P. I., Kawakami, T., Smeds, L., & Ellegren, H. (2013). Demographic Divergence History of Pied Flycatcher and Collared Flycatcher Inferred from Whole-Genome Re-sequencing Data. PLOS Genetics, 9(11), e1003942. https://doi.org/10.1371/journal.pgen.1003942

Nadachowska-Brzyska, K., Burri, R., Smeds, L., & Ellegren, H. (2016). PSMC analysis of effective population sizes in molecular ecology and its application to black-and-white Ficedula flycatchers. Molecular Ecology, 25(5), 1058–1072. https://doi.org/10.1111/mec.13540

Nater, A., Burri, R., Kawakami, T., Smeds, L., & Ellegren, H. (2015). Resolving Evolutionary Relationships in Closely Related Species with Whole-Genome Sequencing Data. Systematic Biology, 64(6), 1000–1017. https://doi.org/10.1093/sysbio/syv045

Nielsen, R., Williamson, S., Kim, Y., Hubisz, M. J., Clark, A. G., & Bustamante, C. (2005). Genomic scans for selective sweeps using SNP data. Genome Research, 15(11), 1566– 1575. https://doi.org/10.1101/gr.4252305

Nosil, P., Funk, D. J., & Ortiz-Barrientos, D. (2009). Divergent selection and heterogeneous genomic divergence. Molecular Ecology, 18(3), 375–402. https://doi.org/10.1111/j.1365-294X.2008.03946.x

Pool, J. E., & Nielsen, R. (2007). Population Size Changes Reshape Genomic Patterns of Diversity. Evolution, 61(12), 3001–3006. https://doi.org/10.1111/j.1558-5646.2007.00238.x

Presgraves, D. C. (2008). Sex chromosomes and speciation in Drosophila. Trends in Genetics, 24(7), 336–343. https://doi.org/10.1016/j.tig.2008.04.007

Presgraves, D. C. (2018). Evaluating genomic signatures of “the large X-effect” during complex speciation. Molecular Ecology, 27(19), 3822–3830. https://doi.org/10.1111/mec.14777

Qvarnström, A., Rice, A. M., & Ellegren, H. (2010). Speciation in Ficedula flycatchers. Philosophical Transactions of the Royal Society B: Biological Sciences, 365(1547), 1841–1852. https://doi.org/10.1098/rstb.2009.0306

Ravinet, M., Faria, R., Butlin, R. K., Galindo, J., Bierne, N., Rafajlović, M., Noor, M. a. F., Mehlig, B., & Westram, A. M. (2017). Interpreting the genomic landscape of speciation: A road map for finding barriers to gene flow. Journal of Evolutionary Biology, 30(8), 1450–1477. https://doi.org/10.1111/jeb.13047

Rice, W. R. (1984). Sex Chromosomes and the Evolution of Sexual Dimorphism. Evolution, 38(4), 735–742. https://doi.org/10.2307/2408385

Sæther, S. A., Sætre, G.-P., Borge, T., Wiley, C., Svedin, N., Andersson, G., Veen, T., Haavie, J., Servedio, M. R., Bureš, S., Král, M., Hjernquist, M. B., Gustafsson, L., Träff, J., & Qvarnström, A. (2007). Sex Chromosome-Linked Species Recognition and Evolution of Reproductive Isolation in Flycatchers. Science, 318(5847), 95–97. https://doi.org/10.1126/science.1141506

Sætre, G., Borge, T., Lindroos, K., Haavie, J., Sheldon, B. C., Primmer, C., & Syvänen, A. (2003). Sex chromosome evolution and speciation in Ficedula flycatchers. Proceedings of the Royal Society of London. Series B: Biological Sciences, 270(1510), 53–59. https://doi.org/10.1098/rspb.2002.2204

Sætre, G.-P., & Sæther, S. A. (2010). Ecology and genetics of speciation in Ficedula flycatchers. Molecular Ecology, 19(6), 1091–1106. https://doi.org/10.1111/j.1365-294X.2010.04568.x

Schumer, M., Xu, C., Powell, D. L., Durvasula, A., Skov, L., Holland, C., Blazier, J. C., Sankararaman, S., Andolfatto, P., Rosenthal, G. G., & Przeworski, M. (2018). Natural selection interacts with recombination to shape the evolution of hybrid genomes. Science, 360(6389), 656–660. https://doi.org/10.1126/science.aar3684

Segami, J. C., Mugal, C. F., Cunha, C., Bergin, C., Schmitz, M., Semon, M., & Qvarnström, A. (2022). The genomic basis of hybrid male sterility in Ficedula flycatchers(p. 2022.09.19.508503). bioRxiv. https://doi.org/10.1101/2022.09.19.508503

Smeds, L., Qvarnström, A., & Ellegren, H. (2016). Direct estimate of the rate of germline mutation in a bird. Genome Research, 26(9), 1211–1218. https://doi.org/10.1101/gr.204669.116

Storchová, L., & Hořák, D. (2018). Life-history characteristics of European birds. Global Ecology and Biogeography, 27(4), 400–406. https://doi.org/10.1111/geb.12709

Storchová, R., Reif, J., & Nachman, M. W. (2010). Female Heterogamety and Speciation: Reduced Introgression of the Z Chromosome Between Two Species of Nightingales. Evolution, 64(2), 456–471. https://doi.org/10.1111/j.1558-5646.2009.00841.x

Svedin, N., Wiley, C., Veen, T., Gustafsson, L., & Qvarnström, A. (2008). Natural and sexual selection against hybrid flycatchers. Proceedings of the Royal Society B: Biological Sciences, 275(1635), 735–744. https://doi.org/10.1098/rspb.2007.0967

Svensson, L., Collinson, J. M., Knox, A., Parkin, D. T., & Sangster, G. (2005). Species limits in the Red-breasted Flycatcher. British Birds, 98, 538–541.

Uebbing, S., Künstner, A., Mäkinen, H., Backström, N., Bolivar, P., Burri, R., Dutoit, L., Mugal, C. F., Nater, A., Aken, B., Flicek, P., Martin, F. J., Searle, S. M. J., & Ellegren, H. (2016). Divergence in gene expression within and between two closely related flycatcher species. Molecular Ecology, 25(9), 2015–2028. https://doi.org/10.1111/mec.13596

Via, S., & West, J. (2008). The genetic mosaic suggests a new role for hitchhiking in ecological speciation. Molecular Ecology, 17(19), 4334–4345. https://doi.org/10.1111/j.1365-294X.2008.03921.x

Vicoso, B., & Charlesworth, B. (2009). Effective Population Size and the Faster-X Effect: An Extended Model. Evolution, 63(9), 2413–2426. https://doi.org/10.1111/j.1558-5646.2009.00719.x

Wang, Z., Zhang, J., Yang, W., An, N., Zhang, P., Zhang, G., & Zhou, Q. (2014). Temporal genomic evolution of bird sex chromosomes. BMC Evolutionary Biology, 14(1), 250. https://doi.org/10.1186/s12862-014-0250-8

Wolf, J. B. W., & Ellegren, H. (2017). Making sense of genomic islands of differentiation in light of speciation. Nature Reviews Genetics, 18(2), Article 2. https://doi.org/10.1038/nrg.2016.133

