## Supplementary Material for "Evidence that genetic drift not adaptation drives *fast-Z* and *large-Z* effects in *Ficedula* flycatchers"

### Supplemental tables

**Table S1:** Sample information of individuals used for PSMC analysis

**Table S2:** Estimates of diversity and PSMC historical population size on autosomes and the Z-chromosome

**Table S3:** Estimates of diversity on autosomes and the Z-chromosome for GC-conservative polymorphisms

**Table S4:** Estimates of selection on protein-coding sequences for autosomes and the Z-chromosome

**Table S5:** Number of nonsynonymous fixed differences overlapping with selective sweeps

**Table S6:** Density of fixed differences in neutral regions and functionally constrained regions on the autosomes vs. the Z-chromosome

**Table S7:** The number of shared SNPs on the autosomes vs. the Z-chromosome for both species comparisons

**Table S8:** The number of outlier $f_d$ windows on the autosomes vs. the Z-chromosome

### Supplemental figures

**Figure S1:** Historical changes in population size for all four species estimated on the Z-chromosome

**Figure S2:** Distributions of $F_{ST}$ estimates across chromosomes

**Figure S3:** Distributions of CLR estimates across chromosomes

**Figure S4:** Distributions of $f_d$ estimates across chromosomes

**Table S1: Sample information of individuals used for PSMC analysis.** Shown are the sample IDs, sex and average sequencing coverage for each individual chosen for the PSMC analysis.

| Species | Sample ID | Sex | Average coverage |
|---|---|---|---|
| **Collared** | Sample_15M58 | Male | 54 |
| **Pied** | SP_14_M | Male | 57 |
| **Red-breasted** | Sample_83 | Male | 65 |
| **Taiga** | Sample_3 | Male | 52 |

**Table S2: Estimates of diversity and PSMC historical population size on autosomes and the Z-chromosome.** Shown are estimates of nucleotide diversity for all four species, estimated using all SNPs on the autosomes and the Z-chromosome separately (see Supplemental Table S3 for GC-conservative estimates). Diversity estimates are also provided after masking sites in exons , conserved non-coding elements and their respective 1kb flanking regions, to account for the impact of linked selection (No LS). PSMC historical population size estimates for all four species are shown for the last 1Mya.

| | **Collared** | **Pied** | **Red-breasted** | **Taiga** |
|---|---|---|---|---|
| $\pi_A$ | 0.00291 [0.00291;0.00292] | 0.00235 [0.00235;0.00236] | 0.00297 [0.00297;0.00297] | 0.00305 [0.00305;0.00306] |
| $\pi_Z$ | 0.00205 [0.00205;0.00206] | 0.00153 [0.00153;0.00154] | 0.00244 [0.00244;0.00245] | 0.00261 [0.00261;0.00262] |
| $\pi_Z/\pi_A$ | 0.705 [0.703;0.706] | 0.651 [0.649;0.653] | 0.822 [0.821;0.825] | 0.855 [0.853;0.858] |
| $\pi_A$ (No LS) | 0.00317 [0.00317;0.00318] | 0.00256 [0.00256;0.00256] | 0.00321 [0.003120;0.00321] | 0.00332 [0.00332;0.00333] |
| $\pi_Z$ (No LS) | 0.00219 [0.00218;0.00219] | 0.00164 [0.00164;0.00165] | 0.00260 [0.00260;0.00261] | 0.00278 [0.00277;0.00278] |
| $\pi_Z/\pi_A$ (No LS) | 0.689 [0.687;0.692] | 0.640 [0.638;0.642] | 0.812 [0.810;0.814] | 0.835 [0.832;0.834] |
| **$PSMC_A$** (until 1Mya) | 182371 [177155, 186660] | 137778 [133726, 142001] | 123549 [120645, 127617] | 96630 [94756, 103304] |
| **$PSMC_Z$** (until 1Mya) | 132586 [127659, 138730] | 92983 [89281, 97015] | 91653 [86859, 95831] | 79408 [69768, 80976] |
| **$PSMC_{Z/A}$** (until 1Mya) | 0.727 [0.691, 0.768] | 0.675 [0.636, 0.707] | 0.742 [0.707, 0.779] | 0.822 [0.706, 0.831] |

**Table S3: Estimates of diversity on autosomes and the Z-chromosome for GC-conservative polymorphisms.** Shown are estimates of nucleotide diversity for all four species, estimated using only GC-conservative SNPs on the autosomes and the Z-chromosome separately. Diversity estimates are also provided after masking sites in exons or conserved non-coding elements, to account for the impact of linked selection.

| | **Collared** | **Pied** | **Red-breasted** | **Taiga** |
|---|---|---|---|---|
| $\pi_A$ | 0.000465 [0.000464;0.000465] | 0.000370 [0.000370;0.000371] | 0.000518 [0.000517;0.000518] | 0.000511 [0.000510;0.000511] |
| $\pi_Z$ | 0.000334 [0.000331;0.000336] | 0.000247 [0.000246;0.000249] | 0.000437 [0.000435;0.000440] | 0.000444 [0.00441;0.000447] |
| $\pi_Z/\pi_A$ | 0.718 [0.713;0.729] | 0.668 [0.663;0.672] | 0.845 [0.840;0.849] | 0.869 [0.864;0.874] |
| $\pi_A$ (No LS) | 0.000511 [0.000510;0.000512] | 0.000407 [0.000406;0.000408] | 0.000564 [0.000563;0.000565] | 0.000562 [0.000561;0.000563] |
| $\pi_Z$ (No LS) | 0.000358 [0.000355;0.000361] | 0.000265 [0.000263;0.000268] | 0.000472 [0.000469;0.000475] | 0.000475 [0.000472;0.000479] |
| $\pi_Z/\pi_A$ (No LS) | 0.700 [0.695;0.706] | 0.652 [0.647;0.657] | 0.837 [0.831;0.842] | 0.845 [0.838;0.851] |

**Table S4: Estimates of selection on protein-coding sequences for autosomes and the Z-chromosome.** Shown are estimates of $\pi_N/\pi_S$, $d_N/d_S$, and $\omega_a$ for genes on the autosomes and on the Z-chromosome. All estimates are based on all sites.

| | **Collared** | **Pied** | **Taiga** | **Red-breasted** |
|---|---|---|---|---|
| **$\pi_N/\pi_S$ A** | 0.186 [0.178;0.194] | 0.197 [0.188;0.207] | 0.175 [0.167;0.183] | 0.175 [0.166;0.182] |
| **$\pi_N/\pi_S$ Z** | 0.201 [0.161;0.237] | 0.287 [0.218;0.333] | 0.209 [0.165;0.270] | 0.178 [0.138;0.213] |
| **$d_N/d_S$ A** | 0.180 [0.173;0.186] | | | |
| **$d_N/d_S$ Z** | 0.191 [0.172;0.209] | | | |
| **$\omega_a$ A** | 0.0506 [0.0423;0.0568] | 0.0370 [0.0237;0.0521] | 0.0903 [0.0822;0.0961] | 0.0287 [0.0200;0.0396] |
| **$\omega_a$ Z** | 0.0752 [0.0369;0.101] | -0.0204 [-0.0591;0.0428] | 0.0729 [0.0402; 0.103] | 0.0823 [0.0357;0.112] |

**Table S5: Number of nonsynonymous fixed differences overlapping with selective sweeps.** Shown for each species are the numbers of nonsynonymous fixed differences identified between the two species comparisons that also show a significant CLR estimate, separately for autosomes and the Z-chromosome. The percent of total nonsynonymous fixed differences showing a sweep signature is given in parentheses for both chromosome types.

| | **Collared** | **Pied** | **Taiga** | **Red-breasted** |
|---|---|---|---|---|
| **Autosomes** | 11 (8.7%) | 41 (32%) | 210 (6.9%) | 394 (11%) |
| **Z-chromosome** | 0 (0%) | 2 (2.4%) | 6 (1.9%) | 3 (0.96%) |

**Table S6: Density of fixed differences in neutral regions and functionally constrained regions on the autosomes vs. the Z-chromosome.** Shown are the proportion of fixed differences for varying functional constraint on autosomes and the Z-chromosome between both collared and pied flycatchers and red-breasted and taiga flycatchers. The fixed differences are separated based on whether they are potentially functionally constrained (UTRs, CNEs, and zero-fold degenerate sites) versus putatively neutrally evolving (intronic regions, intergenic regions, and four-fold degenerate sites). For both species comparisons the results of a Z-test are presented.

| | **Autosome** | **Z-chromosome** | $\chi^2$ | ***P*-value** |
|---|---|---|---|---|
| **Collared and pied constrained** | 0.355 | 0.344 | $2.94 \cdot 10^{-34}$ | 1 |
| **Collared and pied neutral** | 0.645 | 0.656 | | |
| **Red-breasted and taiga constrained** | 0.308 | 0.309 | $5.86 \cdot 10^{-34}$ | 1 |
| **Red-breasted and taiga neutral** | 0.692 | 0.691 | | |

**Table S7: The number of shared SNPs on the autosomes vs. the Z-chromosome for both species comparisons.** In parentheses, the number of shared SNPs are given as a percentage of the total number of sites that are polymorphic in either species within each comparison.

| Chromosome | Collared vs. pied | Red-breasted vs taiga |
|---|---|---|
| **Autosomes** | 2011245 (16%) | 577784 (2.0 %) |
| **Z-chromosome** | 16341 (3.0%) | 16715 (1.3%) |

**Table S8: The number of outlier $f_d$ windows on the autosomes vs. the Z-chromosome.** Shown are the number of windows on the autosomes and the Z-chromosome that have significantly reduced $f_d$ values compared to the genome-wide background for two population comparisons. The population abbreviations are Atlas flycatcher (A), Öland pied flycatcher (OP), Spanish pied flycatcher (EP), Öland collared flycatcher (OC), and Italian collared flycatcher (IC).

| Comparison | Window type | Autosomes | Z-chromosome | Odds ratio | *P*-value |
|---|---|---|---|---|---|
| **((A, OP), OC)** | $f_d$ outlier | 166 | 111 | 0.057 | $< 10^{-16}$ |
| **((A, OP), OC)** | No outlier | 4351 | 165 | | |
| **((A, EP), IC)** | $f_d$ outlier | 159 | 126 | 0.044 | $< 10^{-16}$ |
| **((A, EP), IC)** | No outlier | 4328 | 150 | | |

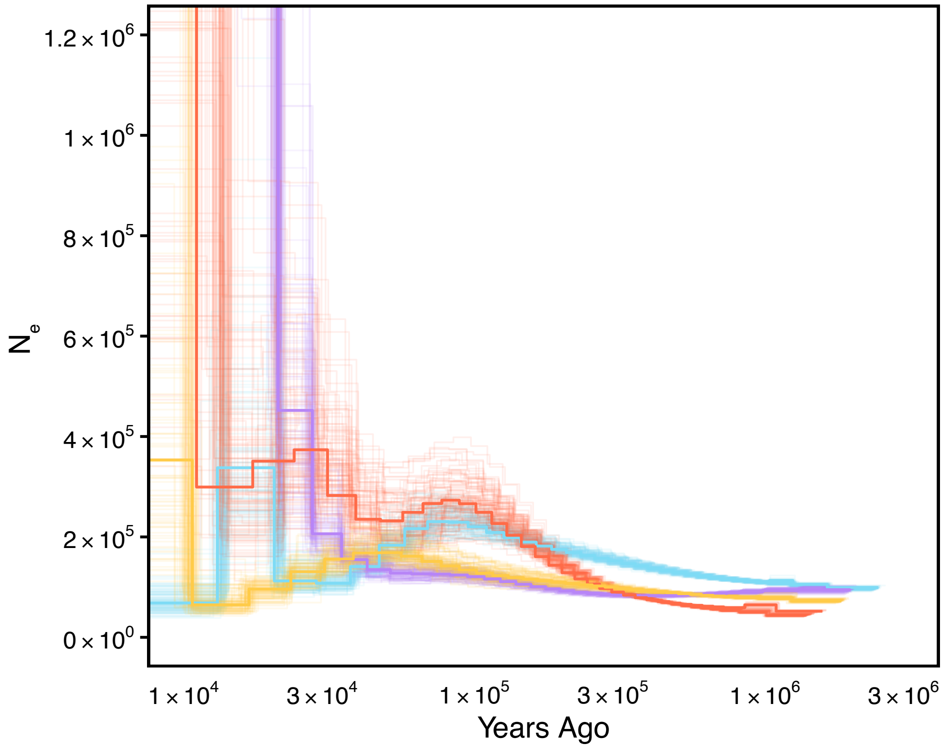


**Figure S1: Historical changes in population size for all four species estimated on the Z-chromosome.** One individual for each of the four flycatcher species is represented: collared flycatcher in blue, pied flycatcher in purple, red-breasted flycatcher in yellow, and taiga flycatcher in orange. The bold line represents the genome-wide estimate, bootstrap replicates are shown in lighter colour.

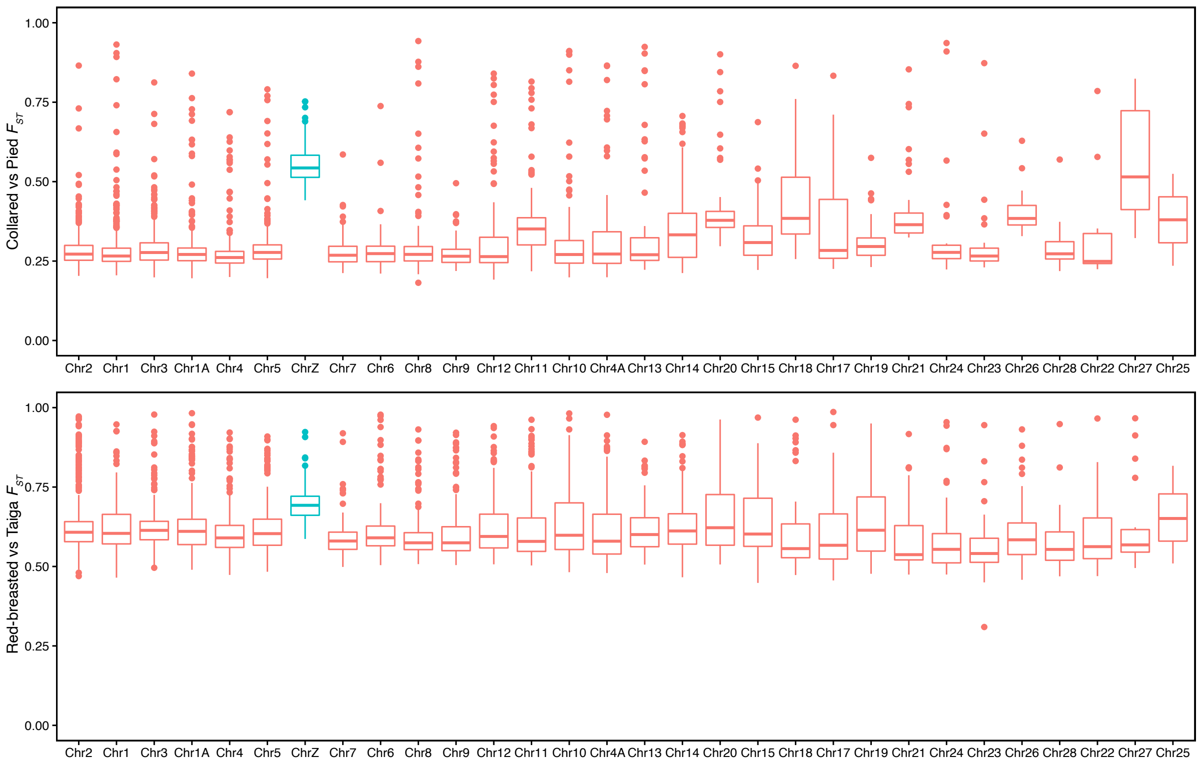


**Figure S2: Distributions of $F_{ST}$ estimates across chromosomes.** Shown are estimates of $F_{ST}$ in 200kb genomic windows for the collared and pied flycatchers and for the red-breasted and taiga flycatchers divided by chromosome. Chromosomes are ordered by decreasing size, with autosomes in red and the Z-chromosome highlighted in blue.

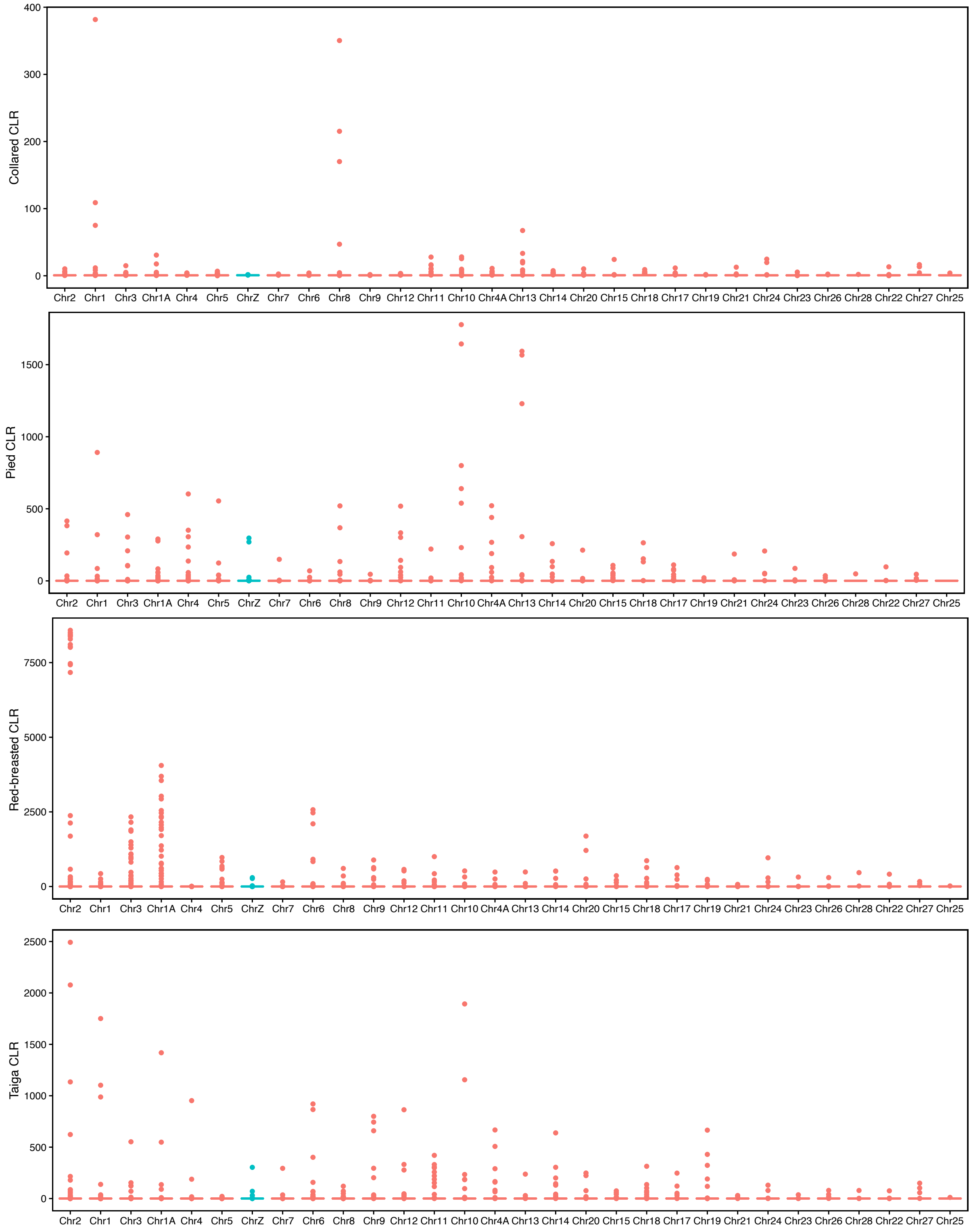


**Figure S3: Distributions of CLR estimates across chromosomes.** Shown are average CLR values in 200kb genomic windows for the four species, collared flycatcher, pied flycatcher, red-breasted flycatcher, and taiga flycatcher divided by chromosome. Chromosomes are ordered by decreasing size, with autosomes in red and the Z-chromosome highlighted in blue.

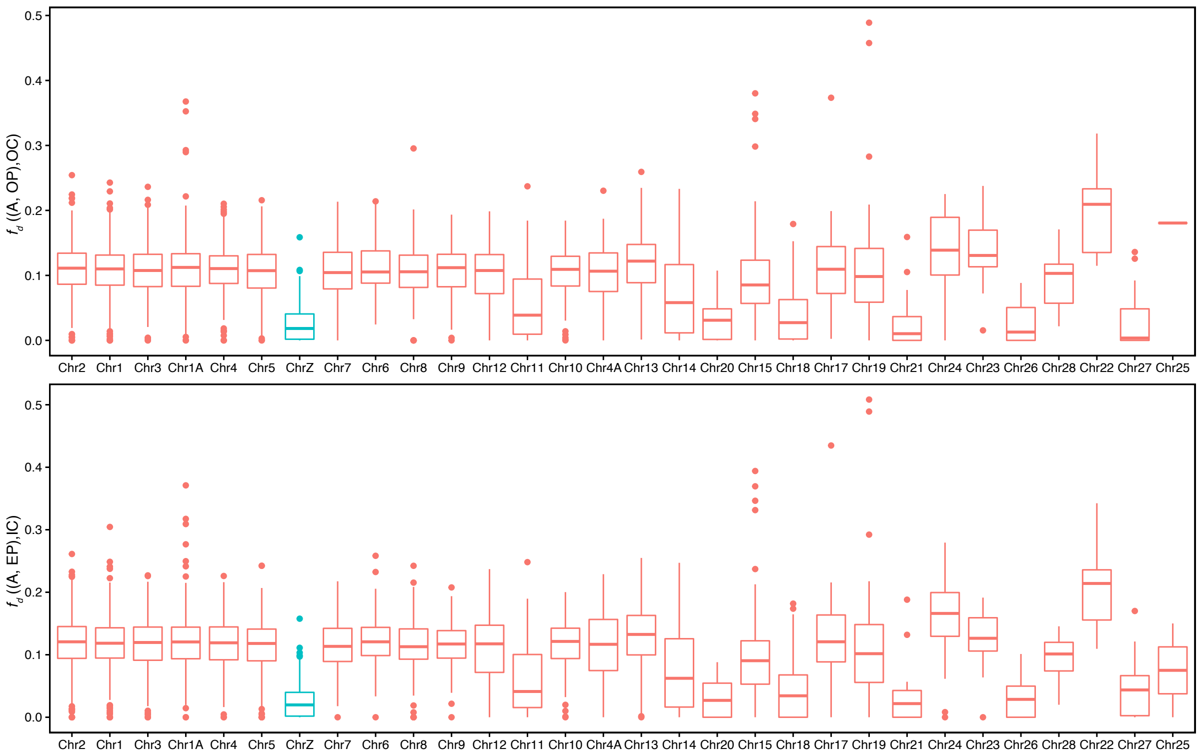


**Figure S4: Distributions of $f_d$ estimates across chromosomes.** Shown are estimates of the $f_d$ statistic in 200kb genomic windows for two population comparisons, divided by chromosome. The population abbreviations are: Atlas flycatcher (A), Öland pied flycatcher (OP), Öland collared flycatcher (OC), Spanish pied flycatcher (EP), and Italian collared flycatcher (IC). Chromosomes are ordered by decreasing size, with autosomes in red and the Z-chromosome highlighted in blue.
